# Comparing Expression of OAS-RNaseL Pathway-Related Genes in SARS-CoV-2 and Similar Viruses

**DOI:** 10.1101/2021.04.29.442063

**Authors:** Nikhil Chakravarty, Penelope A. Edillor, Andy P. Huang, Giovanni A. Torres, Manal T. Usmani, Pavan Kadandale

**Author notes:** NC, PAE, APH, GAT, and MTU all contributed equally to this paper and should be regarded as co-first authors.

## Abstract

The COVID-19 pandemic, caused by the virus SARS-CoV-2, has been a major public health emergency and has caused millions of deaths worldwide to date. Due to the novel nature of the virus, efforts across the world are underway to better understand the molecular pathogenesis of SARS-CoV-2 and how it interacts with host immune responses. One important branch of the innate immune response, the interferon system, triggers the expression of many effector mechanisms known to be powerful antagonists against many pathogenic viruses. One such interferon stimulated mechanism is the OAS-RNaseL pathway, which is known to trigger the degradation of viral RNA in infected host cells. Our study seeks to utilize publicly available transcriptomic data to analyze the host cell OAS-RNaseL pathway to SARS-CoV-2 infection. We hoped to gain an understanding of the importance of the pathway in controlling SARS-CoV-2 infection and whether or not the pathway could be exploited therapeutically. Our findings demonstrated that upregulation of OAS-RNaseL pathway genes in response to SARS-CoV-2 infection varies based on cell type and appeared to correlate with ACE2 receptor expression. Pathway responses to other viruses like SARS-CoV and MERS-CoV were found to parallel those to SARS-CoV-2, suggesting common response patterns by the pathway to these viruses. Overall, these results demonstrate that the OAS-RNaseL pathway could contribute to control of SARS-CoV-2 infection. Further studies on various mechanistic actions by the pathway would need to be conducted to fully understand its role in host defense and therapy.

## Introduction

Since being deemed a pandemic in March 2020, coronavirus disease 2019 (COVID-19) has taken the lives of millions worldwide as SARS-CoV-2, its causative agent, has infected tens of millions at the time of publication^1^. The novel coronavirus SARS-CoV-2, a positive-sense, single-stranded RNA virus, is part of the *Coronaviridae* family and *Orthocoronavirinae* subfamily^2^. MERS-CoV and SARS-CoV, two other members of the *Orthocoronavirinae* subfamily, were responsible for past epidemics and are highly pathogenic to humans, whereas the other four are only mildly pathogenic^3,4^. SARS-CoV-2, SARS-CoV, and MERS-CoV cause common clinical symptoms including cough, congestion, fatigue, shortness of breath, and fever, that all contribute to acute respiratory distress syndrome (ARDS)^5^. However, SARS-CoV-2 infection is unique in that these symptoms can be highly variable in severity. In addition, the SARS-CoV-2 virus has a longer incubation period, facilitating greater asymptomatic transmission to a degree not seen with SARS-CoV and MERS-CoV^6^. These characteristics, as well as the novel nature of the SARS-CoV-2 virus, justify the need to further investigate the various immune responses to a SARS-CoV-2 infection and how such responses compare to those generated by similar viruses.

A major mechanism of the human innate immune system activated against viral infection is the interferon (IFN) signaling system. In human cells, a variety of pattern recognition receptors (PRRs) recognize pathogen-associated molecular patterns (PAMPs) and induce signaling cascades that lead to interferon production^7^. In the context of RNA viruses like coronaviruses, several toll-like receptors (TLRs) and RIG-I-like receptors (RLRs) recognize double-stranded RNA (dsRNA) as a PAMP and signal to primarily induce type I and type III interferon expression^7^. Type I interferons, the most well understood category, engage the IFNAR receptor via autocrine or paracrine signaling and induce a signaling cascade leading to the production of interferon-stimulated genes (ISGs)^7,8^. One particular group of ISGs are the oligoadenylate synthetases (OAS) – proteins that have the ability to catalyze the polymerization of ATP in a 2’-5’ fashion to create 2’-5’ oligoadenylate (2-5A)^8^ (**Figure 1**). 2-5A activates the latent RNaseL, leading to the degradation of cytosolic RNA and subsequent disruption of viral replication^9^. This process is facilitated by the destruction of ssRNA viral genomes and mRNA used for viral gene expression, the production of small duplex RNAs leading to further IFN-β production, and the degradation of rRNA and critical components of host cell machinery leading to apoptosis^9^. The OAS-RNaseL pathway has been found to possess antiviral activity against a wide variety of RNA and DNA viruses, including Respiratory Syncytial Virus (RSV), West Nile Virus, Hepatitis C Virus, Human Immunodeficiency Virus type I, and Herpes Simplex Virus type I (HSV-1)^10–13,16^. However, many viruses, including MERS-CoV, Influenza A virus (IAV), and Human Parainfluenza Virus 3 (HPIV3), have established mechanisms to inhibit the activity of this pathway^14,15,17^. For example, the NS4b protein of MERS-CoV functions as a viral phosphodiesterase which degrades 2’-5’ oligoadenylate^14^. Several host proteins have also been found to antagonize various aspects of the OAS-RNaseL pathway, such as the mammalian protein ABCE1, an RNaseL inhibitor, and the AKAP7, PDE12, and ENPP1 proteins, which all possess a phosphodiesterase function which allow them to degrade 2-5A^18–21^.

**Figure 1.**
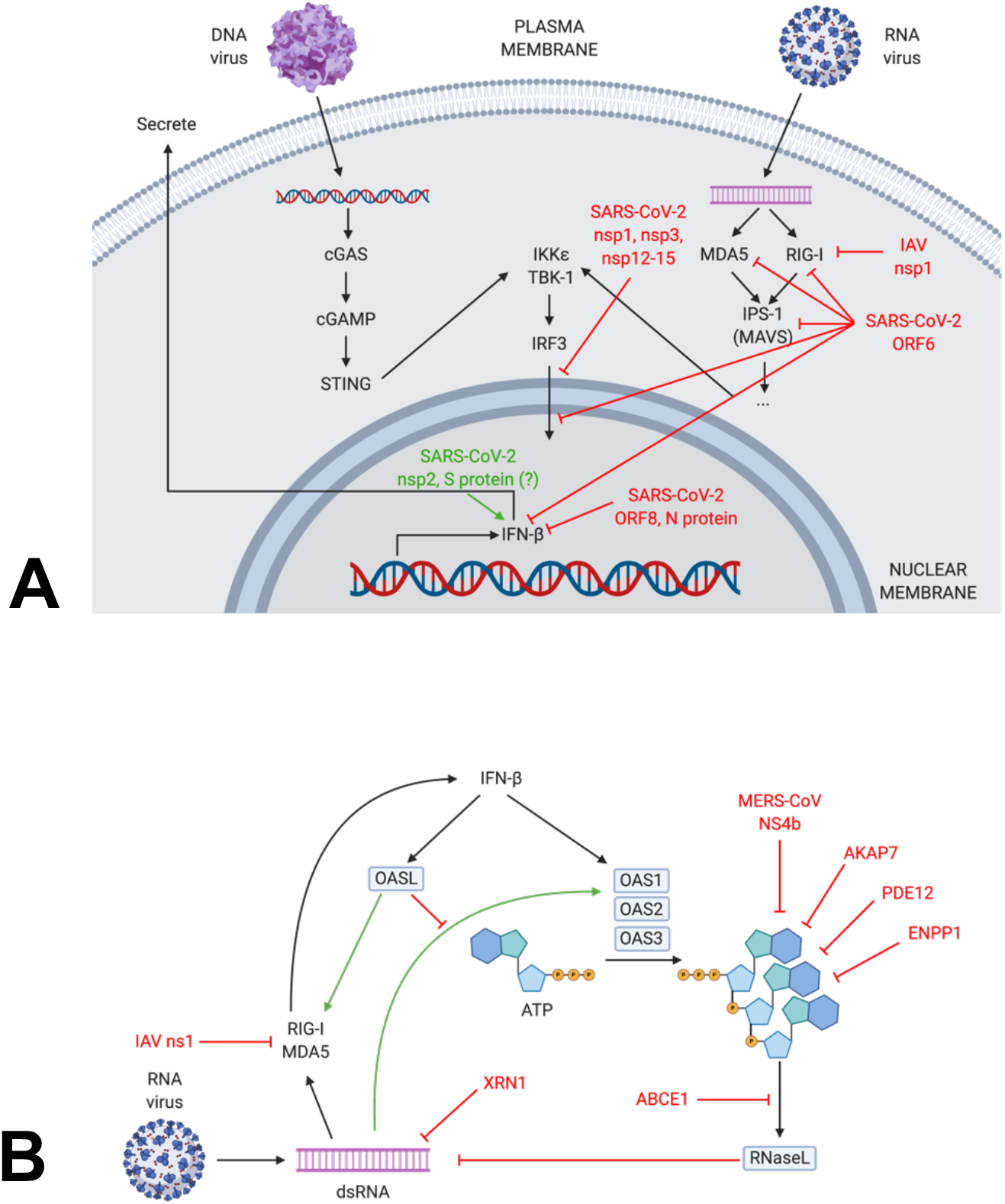
Type I Interferon and OAS-RNaseL innate immune pathways. Type I interferon signaling induced by RNA and DNA viruses (A) and antiviral activity of 2’-5’ oligoadenylate synthetase and RNaseL (OAS-RNaseL) pathway induced by IFN-β (B)^7–21^. Figures created with BioRender.com.

IFN signaling pathways are known to be activated upon SARS-CoV, MERS-CoV, and SARS-CoV-2 infection to modulate the adaptive antiviral response of infected cells^22–24^. For example, IFN-β therapy has been demonstrated to be effective at controlling MERS-CoV infection in marmosets^25^. Due to the similarity between the viruses, IFN therapy has also been discussed as a potential treatment for SARS-CoV and SARS-CoV-2. However, since IFN signaling is itself involved in facilitating cytokine storming, many researchers are skeptical about its effectiveness in severe cases^26^. Another limitation is IFN therapy would need to be administered in the early stages of infection - before symptoms appear^26^. The presence of anti-IFN proteins in a variety of viruses further complicates the potential success of IFN therapy. Type I IFN induction has been shown to vary by cell type and virus type. For example, Blanco-Melo *et al.* infected normal human bronchial epithelial (NHBE) cells with both wild type IAV and a NS1 knockout strain to investigate the role of NS1 in IFN induction^27^. Their results showed that the knockout elicited significantly stronger type I and III interferon responses, confirming the role of viral interferon suppressors like NS1 in NHBE cells^27^. Conversely, the OAS-RNaseL pathway is also exploited therapeutically in a fashion independent of IFN signaling. This pathway can be utilized through two mechanisms: the established RNaseL mechanism and, interestingly, an RNaseL independent mechanism^28^. Kristiansen *et al.* showed that OAS1 can act as a direct, exogenous antiviral compound against Encephalomyocarditis virus, Vesicular Stomatitis Virus (VSV), and HSV-1^28^. This provides a promising result: exogenous OAS could be used as an antiviral agent. As very few, if any, research teams have studied the potential link between the OAS-RNaseL pathway and SARS-CoV-2 pathogenesis, we would like to investigate the expression and role of the OAS pathway in the context of SARS-CoV-2 pathology and compare infections of related viruses in order to suggest further studies using OAS as a treatment for COVID-19 and similar diseases.

To compare the role of the OAS-RNaseL pathway in SARS-CoV-2 infection and those of related viruses, we studied the differences in gene regulation of OAS and IFN-related genes in infected and uninfected cell lines. This comparison was achieved through the analysis of RNAseq data using the online bioinformatics platform, Galaxy. This allowed us to visualize similarities and differences in OAS-RNaseL pathway expression changes between cases of SARS-CoV-2 infection and infection with other types of viruses, including related coronaviruses on top of a variety of other virus types. Upon SARS-CoV-2 infection, we found that OAS proteins and IFN-β were significantly upregulated in cell lines expressing the SARS-CoV-2 entry receptor ACE2. The OAS-RNaseL pathway was also upregulated upon infection with different viruses known to trigger the pathway. Overall, through the comparative nature of this study, we aimed to uncover potential similarities in immune evasion by similar viruses. By determining connections to viruses that already have established therapies, this could bring to light potential therapeutic options to use when treating patients with COVID-19.

## Methods

### RNA sequencing data collection

The NCBI BioProject database (https://www.ncbi.nlm.nih.gov/bioproject/) was searched for RNA-sequencing datasets from experiments with cell cultures or samples infected with SARS-CoV-2 or other viruses between June and November 2020. **Supplemental Table 1** lists the BioProjects and the associated cell lines selected for further analysis, as well as their Sequence Read Archive (SRA) run accessions.

We selected datasets based on various cell line or tissue cultures infected with SARS-CoV-2, related coronaviruses like SARS-CoV and MERS-CoV, or other miscellaneous viruses, including IAV and RSV, using RNAseq technology. The intended goal is to compare gene expression between different tissue cultures infected with SARS-CoV-2 and compare those with other various virus-infected tissue cultures, including different coronaviruses.

Since the goal of this study was to rationalize the relationship between viral infection and stimulation of the OAS-RNaseL pathway, we analyzed the expression levels of genes associated with this pathway.

### RNA sequencing data analysis

The SRA files were extracted by fastq-dump utility of an SRA Toolkit, FASTQ Sequencing, file was uploaded to Galaxy web platform (usegalaxy.org), which was used to perform further data analysis^29^. FastQC Read Quality reports were first generated in Galaxy for each dataset to assess the quality of the data. Next, the Trimmomatic function with the singleend read setting was used to remove Illumina sequencing adapters, remove sequence ends with average quality scores lower than 20 over four bases using sliding window trimming, and remove any sequences smaller than 25 base pairs. New FastQC Read Quality reports were generated after trimming to confirm that necessary changes were made and that sequence quality was acceptable. Comparison between the original FastQC and the Trimmed FastQC reports were conducted to analyze the changes in the total number of sequences and the sequence lengths.

The HISAT2 function was used for each trimmed output to align sequences in each dataset to the genome of their respective organisms. For human datasets, the hg38 genome was used as a reference. Alignment summaries were generated following each HISAT2 analysis to check the percent of sequences successfully aligned to the genome. The featurecounts function, using both built-in and downloaded gene annotation files, was used for each binary alignment map (BAM) output from HISAT2 to count the number of sequences aligned to each gene.

To observe changes in gene expression under different experimental conditions, the DESeq2 function was used to compare counts tables from mock treated datasets and experimentally treated datasets. Genes with a Benjamini-Hochberg p-value of ≤0.01 and log_2_FC≥1 or log_2_FC≤-1 were considered to be significantly up or downregulated, respectively. To determine ACE2 expression levels in different conditions, base means for the ACE2 gene were extrapolated and compared. A base mean of 100 or greater was considered to be significant. For the purposes of this study, genes relating to the OAS-RNaseL pathway, interferon signaling pathways, various viral entry receptors, and a housekeeping gene (β-actin) were analyzed. The respective geneIDs for genes of interest were obtained from the NCBI Gene portal. The results file output from DESeq2 was then searched for these geneIDs to obtain the base mean, log_2_FC values, p-adj values, and other relevant data. The overall workflow through Galaxy is shown in **Figure 2**.

**Figure 2.**
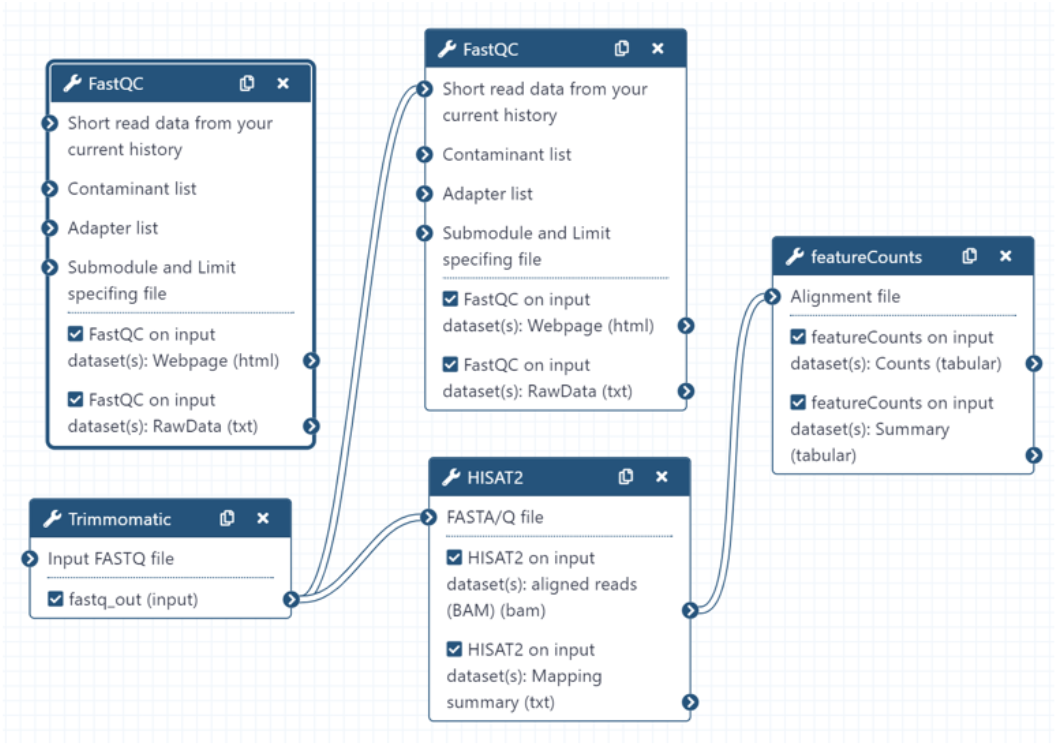
Workflow-construct for RNAseq analysis. The figure illustrates the pipeline used in Galaxy to obtain the appropriate data from specific SRA files. The program allows for an automatic run once files are input (SRR files) and generate output files. While those output files are used for the downstream of generating the wanted featureCounts files.

### Viral Replication Data Collection

The efficacy of the OAS pathway in inhibiting viral replication was assessed across various SARS-CoV-2 infected cell lines, as seen in **Table 1**. The DESeq2 function in Galaxy was used to obtain the ACE2 Base Means and the log_2_FC values of the OAS genes in these cells. The ACE2 Base Means quantified the abundance of the ACE2 receptors. ACE2 receptor abundance was categorized as high if the base mean was greater than 100 and was categorized as low if it was less. The log_2_FC values quantified the strength of OAS gene expression. Log_2_FC values greater than 4 indicated strong OAS gene upregulation, log_2_FC values between 4 and 1.5 indicated moderate upregulation, log_2_FC values between 1.5 and 0 indicated weak upregulation, while log_2_FC values less than or equal to 0 indicated insignificant gene expression. The infection efficiency in these cells was quantified by the percent mapping rate of the intracellular viral RNAs to the viral genome. These values were obtained from the data published by Cao et. al and Sharma *et. al*^30,31^. Percent mapping rates greater than 50% represented medium infection efficiency, rates between 50% and 0% represented low infection efficiency, and rates less than or equal to 0% indicated that there was no viral infection.

**Table 1.**
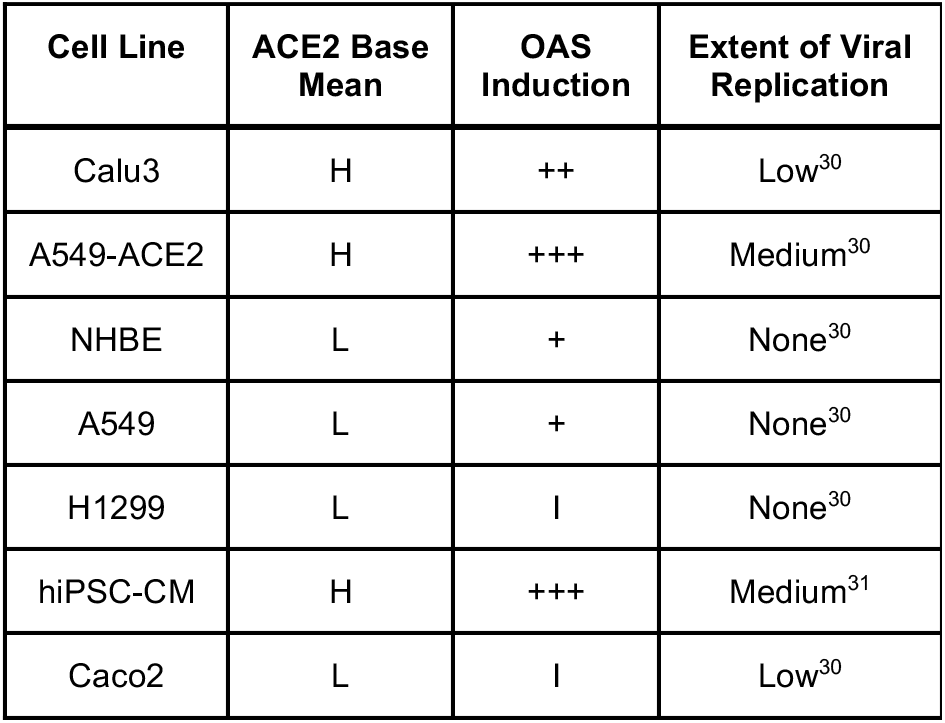
ACE2 base means, OAS induction level, and extent of viral replication for studied cell lines. Experimental data for ACE2 base means and OAS gene induction levels for representative cell lines analyzed in this study along with SARS-CoV-2 replication levels from literature, were collected for comparison.^30,31^ ACE2 base means were categorized as high if levels were greater than 100 and as low if less than 100. OAS induction levels were categorized based on average log_2_FC for OAS1, OAS2, OAS3, and OASL as follows: +++: log_2_FC > 4, ++: 4 ≥ log_2_FC > 1.5, +: 1.5 ≥ log_2_FC > 0. SARS-CoV-2 replication data was based on RNASeq mapping rates to the SARS-CoV-2 genome and was categorized based on the following criteria: medium: % mapped > 50%, low: 50% > % mapped > 0%, none: % mapped ≤ 0%.

## Results

### Expression Profile of OAS-RNaseL Pathway Genes Across Different Cell Lines and Tissues

log_2_FC values of various OAS-RNaseL pathway genes from different cell lines and patient samples were obtained from RNAseq data available from BioProjects and assembled into heat maps. We first compared gene expression in different cell lines and tissues following SARS-CoV-2 infection (**Figure 3**). Calu3, A549-ACE2 infected with a SARS-CoV-2 MOI of 2, hiPSC-CM, HUVEC, and HT-29 displayed strong upregulation of all four *OAS* and *IFNB1* genes simultaneously after SARS-CoV-2 infection (24hpi for Calu3 and A549-ACE2, 72hpi for hiPSC-CM, and 3dpi for HUVEC and HT-29). However, insignificant changes in RNaseL expression were observed. Of these cell lines, HT-29 and HUVEC also expressed the greatest downregulation of genes that inhibit the OAS-RNaseL pathway. Calu3 displayed the greatest upregulation of *IFNB1* and other type I interferons. NHBE, A549 (without a human ACE2 vector and infected with a viral MOI of 0.2), PBMCs expressing high ACE2 base means from COVID-19 patients, and hBEpC lung organoids displayed slight but significant upregulation of OAS genes but insignificant changes in expression of all type I IFN and RNaseL genes (24hpi for NHBE and A549). These cell lines also did not experience significant changes in expression of OAS-RNaseL pathway-inhibiting genes. Lung biopsies from COVID-19 patients exhibited downregulation of OAS genes and very slight expression of type I IFN and RNAseL genes. PBMCs with low ACE2 base means and platelets from COVID-19 patients under intensive care displayed insignificant changes in type I IFN, OAS, and RNaseL expression and exhibited slight and insignificant changes in OAS-RNaseL pathway inhibiting genes. H1299, pHAE, and Caco2 showed no significant changes in expression of type I IFN, OAS, RNaseL or OAS-RNaseL pathway inhibiting genes (24hpi for H1299 and Caco2, 48hpi for pHAE).

**Figure 3.**
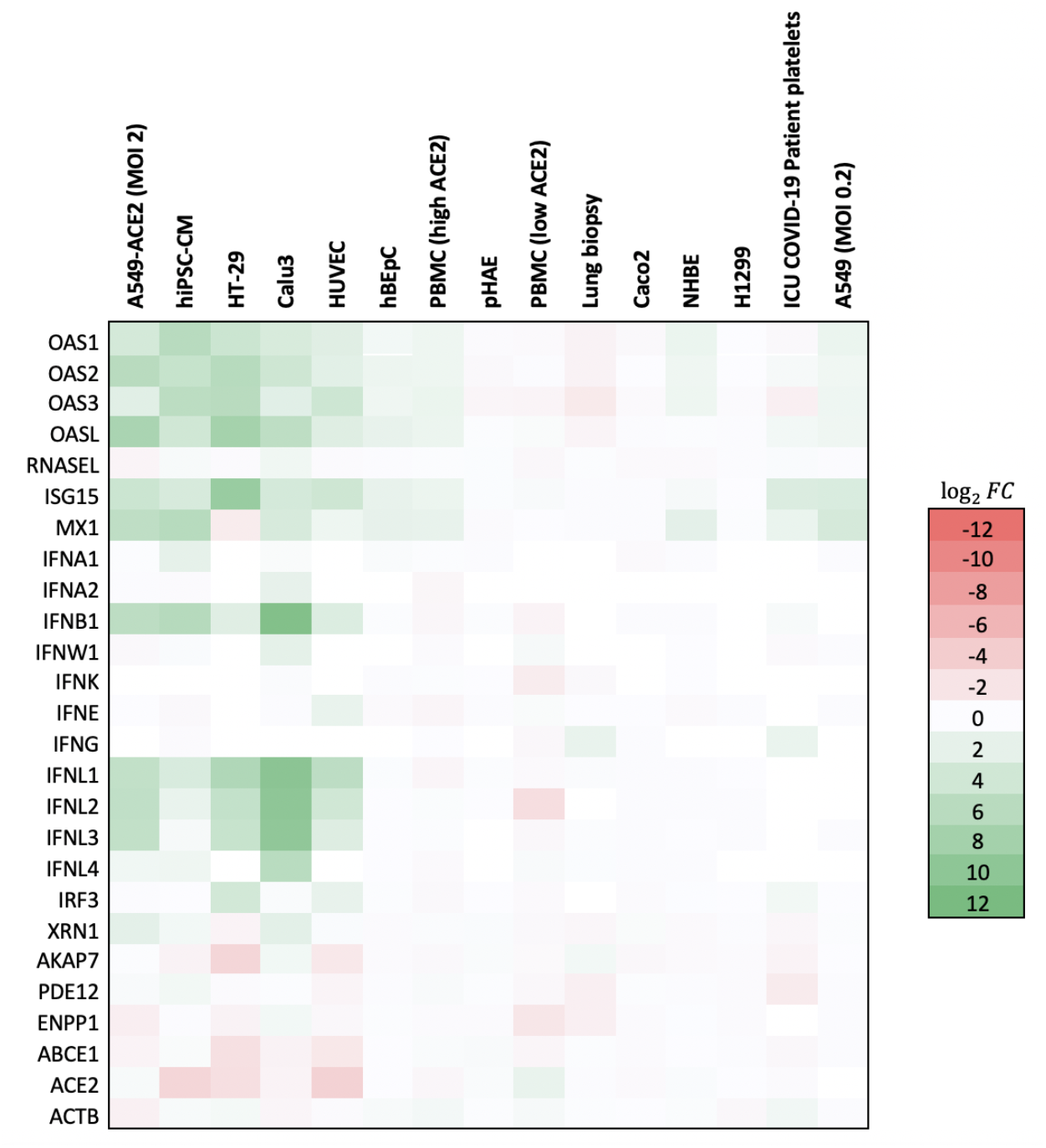
OAS-RNaseL pathway induction in SARS-CoV-2 infection of different tissues. Fold changes in expression of various genes associated with the OAS/RNaseL antiviral pathway in different tissues after SARS-CoV-2 infection. All four OAS genes, along with *IFNB1*, were significantly upregulated in Calu3, A549-ACE2 with a viral MOI of 2, hiPSC-CM, HT-29, and HUVEC. OAS genes were also significantly upregulated in NHBE and A549 with a viral MOI of 0.2, and PBMCs with high ACE2 base means. Conditions are arranged based on ACE2 base mean.

### Quantifying ACE2 Abundance Across Different Cell Lines and Tissues

To explore whether or not SARS-CoV-2 entry receptor abundance correlates with type I IFN and OAS-RNaseL pathway induction, base mean values of the ACE2 gene from different cell lines and patient samples were obtained to quantify expression of the ACE2 receptor (**Figure 4**). A549-ACE2 displayed the highest ACE2 base mean and thus the greatest abundance of the ACE2 gene. Other samples that displayed high ACE2 base means were Calu3, hiPSC-CM, HT-29, and HUVEC. The latter four, along with A549-ACE2, also displayed the greatest simultaneous upregulation of all four OAS genes and IFNB1 (**Figure 3**). Platelets from COVID-19 patients under intensive care displayed low ACE2 base means while the A549 cell line displayed an ACE2 base mean of zero, indicating a low abundance of ACE2 genes in these samples (**Figure 4**). Samples with low ACE2 base means exhibited mostly insignificant changes in OAS-RNaseL and type I IFN expression, with only a handful showing significant upregulation of OAS genes (PBMCs with high ACE2 base mean, NHBE, and A549 without ACE2 vector). This suggests that entry receptor expression could relate to the ability of the virus to induce type I IFN and ISG expression.

**Figure 4.**
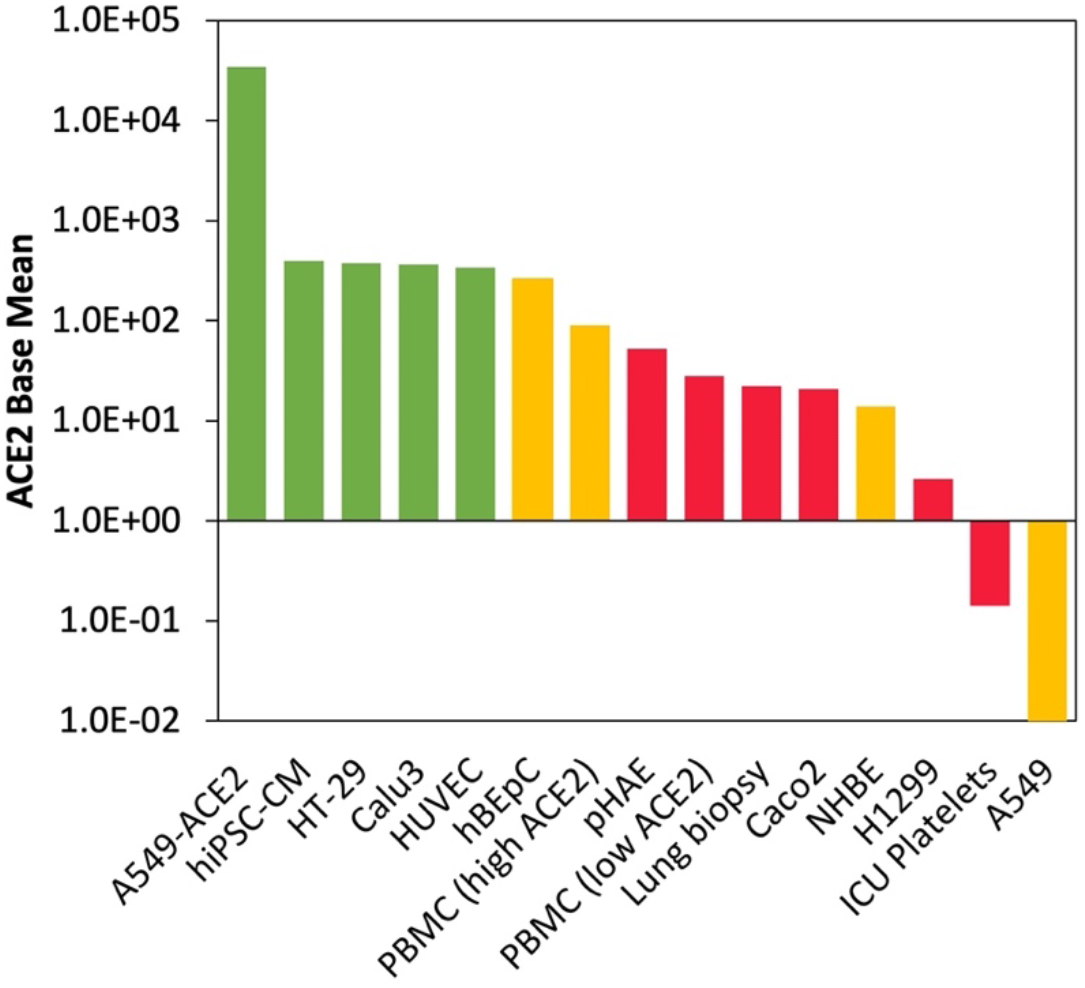
ACE2 normalized counts in different cell lines and tissues. Base means for Angiotensin-converting enzyme 2 (*ACE2*, GeneID: 59272), the entry receptor used by SARS-CoV-2 and several other coronaviruses, in different cell lines, tissues, and biological samples as determined using DESeq2. ACE2 expression appears to vary widely between different tissues and cell lines. A549 cells transduced with an ACE2 vector, unsurprisingly, displayed the highest base mean for ACE2, while Calu3, hiPSC-CM, HT-29, and HUVEC also displayed high ACE2 base means. Different tissues are categorized based on OAS-RNaseL pathway and *IFNB1* induction levels observed following SARS-CoV-2 infection. Green bars indicate that the cell line or sample was found to induce high levels of IFN-β and OAS gene expression. Yellow bars indicate that the sample was found to induce moderate levels of OAS but no interferon gene expression. Red bars indicate that the sample failed to induce significant levels of interferon and OAS expression.

### Comparison of OAS-RNaseL Pathway Induction Across Different Coronaviruses

OAS-RNaseL pathway gene expression between SARS-CoV-2, SARS-CoV, and MERS-CoV was compared by creating an expression profile using the log_2_FC values from the RNAseq analysis of infected Calu3, H1299, and Caco2 cells to better understand the similarities and differences in OAS-RNaseL pathway induction in different organ cells for these viruses. For all three cell lines, infection with SARS-CoV-2 and SARS-CoV displayed very similar expression profiles for type I IFN and OAS-RNaseL pathway genes. In Calu3, both SARS-CoV-2 and SARS-CoV infected cells displayed strong upregulations of all *OAS* genes and *IFNB1*, along with insignificant changes for RNaseL and OAS pathway inhibiting genes **(Figure 5A)**. IFNA2 and IFNW1 were also significantly upregulated in SARS-CoV-2-infected Calu3 cells but not for SARS-CoV- or MERS-CoV-infected cells. MERS-CoV-infected Calu3 cells displayed a similar pattern with slightly weaker OAS gene upregulation and significantly weaker *IFNB1* gene upregulation. H1299 and Caco2 cells infected with SARS-CoV and SARS-CoV-2 displayed little to no changes in OAS-RNaseL pathway gene expression (**Figure 5B, 5C**).

**Figure 5.**
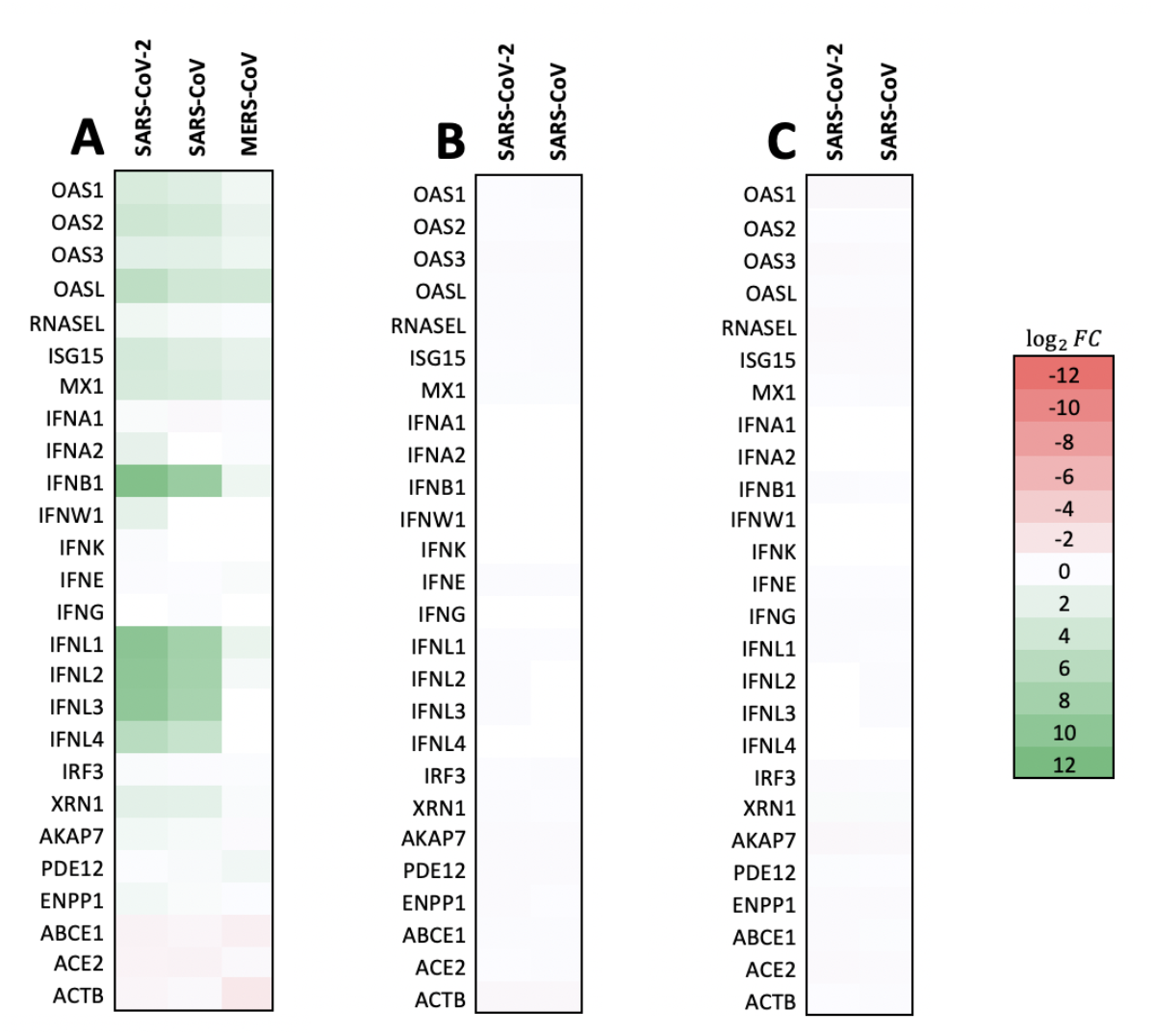
OAS-RNaseL Pathway Induction for Coronaviruses in Lung and Colorectal Cell Cultures. Fold changes in expression of various genes associated with the OAS/RNaseL antiviral pathway in (A) Calu3, (B) H1299, and (C) Caco2 after 24 hours of infection with various coronaviruses. SARS-CoV-2 and SARS-CoV elicited similar patterns of OAS upregulation. OAS expression in Calu3 infected with MERS-CoV followed a similar pattern but genes of interest were not upregulated as heavily. SARS-CoV-2 and SARS-CoV infection of H1299 and Caco2 did not significantly alter OAS pathway gene expression.

### Comparison of OAS-RNaseL Pathway Induction Across Non-Coronaviruses

Similar to conditions with coronaviruses, we compared the OAS-RNaseL pathway gene expression profile of SARS-CoV-2 infected cell lines to those infected with other viruses. Specifically, RNAseq data from A549 cells infected with SARS-CoV-2, RSV, IAV, and HPIV3 were analyzed. A549 cells infected with SARS-CoV-2 at an MOI of 0.2 displayed moderate but significant upregulation of all OAS genes but insignificant changes in expression of type I IFNs, RNaseL, and OAS-RNasL pathway inhibiting genes **(Figure 6A)**. In comparison, A549 infected with RSV (MOI 15) and HPIV3 (MOI 3) also lacked significant changes in expression of RNaseL and OAS-pathway inhibiting genes but exhibited significantly greater upregulation of OAS genes 24-hpi. These two cell lines also displayed low expression of most type I IFN genes, except for *IFNB1*, with HPIV3-infected cells displaying significantly greater upregulation of *IFNB1* compared to RSV-infected cells. HPIV3’s expression profile is similar to that of SARS-CoV-2, SARS-CoV, and MERS-CoV infected Calu3 cells. No significant induction of OAS-RNaseL pathway genes and type I IFN genes were observed in IAV-infected A549 cells 9-hpi. In fact, OAS2 and IRF3 were significantly downregulated whereas AKAP7 and ABCE1 were significantly upregulated in this A549 line **(Figure 6A)**.

**Figure 6.**
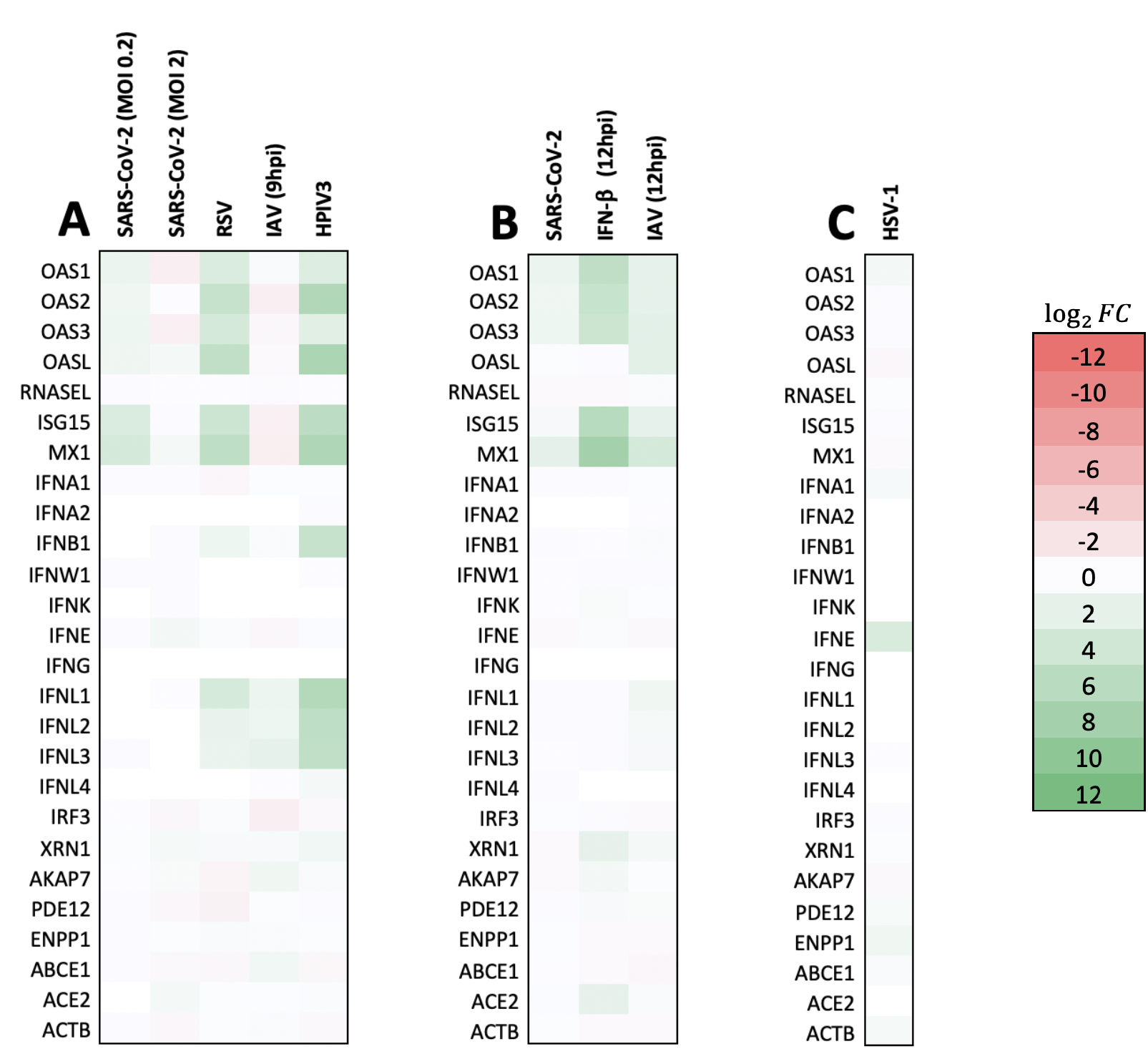
OAS-RNaseL pathway induction for miscellaneous viruses and IFN-β treatment in lung epithelial and fibroblast cells. Fold changes in expression of various genes associated with the OAS/RNaseL antiviral pathway in A549 (A) or NHBE (B) after infection with various viruses or IFN-β treatment. Differential expression for Herpes simplex virus-1 (HSV-1) was also analyzed in normal human dermal fibroblast (NHDF) cells (C). Infection of A549 with SARS-CoV-2 of 0.2 MOI resulted in modest but significant upregulation of OAS, while RSV and HPIV3 resulted in higher levels of OAS upregulation. IAV (which has a known type I interferon antagonizing protein) infection of A549 resulted in no upregulation of OAS at 9hpi (which contradicts prior literature). In NHBE, both SARS-CoV-2 and IAV elicited weaker OAS upregulation compared to IFN-β treatment. HSV-1 elicits a different expression pattern in NHDF cells: IFNE upregulation predominates, but OAS upregulation is insignificant.

NHBE cells infected with SARS-CoV-2 displayed a similar gene expression profile to SARS-CoV-2 infected A549 cells. Both cell lines displayed slight upregulation of OAS genes (except OASL in NHBE) but low inductions of type I IFNs, RNaseL, and OAS-RNaseL pathway inhibiting genes. IAV infected NHBE cells also displayed similar trends, with OASL also being upregulated. IFN-β treated NHBE cells exhibited significantly greater OAS gene upregulation compared to the SARS-CoV-2 and IAV infected NHBE cells (**Figure 6B**).

In addition to lung epithelial cell lines, the OAS-RNaseL pathway expression profile was also assessed for HSV-1, a DNA virus, in NHBE cells. Low expression of most OAS-RNaseL pathway genes but moderate upregulation of the *IFNE* gene was observed. These results are different from the profiles exhibited by RNA viruses in other tissues (**Figure 6C**).

### Relationship Between OAS-RNaseL Induction and SARS-CoV-2 Replication

Given our findings on the activation of the OAS-RNaseL pathway in various cell lines in response to SARS-CoV-2 infection, we wanted to better understand how induction of the pathway correlates to outcomes of infection like the ability for SARS-CoV-2 to proliferate. We compared induction levels of the OAS-RNaseL pathway to data on the replication levels of SARS-CoV-2 in equivalent cells sourced from Cao *et al.* and Sharma *et al.* for hiPSC-CM^30,31^. We found that Calu3, which induces the OAS-RNaseL pathway at a moderately high level, supported low but detectable levels of SARS-CoV-2 replication as detected through RNAseq mapping rates to the SARS-CoV-2 genome **(Table 1)**. However, cells like A549-ACE2 and hiPSC-CM, which induced the pathway at higher levels than Calu3, supported higher levels of SARS-CoV-2 replication. Conversely, NHBE induced the pathway at a lower level but also supported lower levels of viral replication. On the other hand, Caco2 did not significantly induce the pathway but supported detectable levels of SARS-CoV-2 replication. From such a comparison, a direct correlation between OAS-RNaseL induction level and SARS-CoV-2 replication was not apparent. However, given the lack of controlled experimentation regarding this subject, further studies will need to be conducted to fully understand the relationship between OAS, RNaseL, and SARS-CoV-2 replicability.

## Discussion

### OAS Pathway Expression Patterns Differ Across Cell Lines and Could Correlate with ACE2 Abundance

Our results show that a characteristic pattern of *IFNB1* and *OAS* gene upregulation is generally observed when SARS-CoV-2 infects certain cell types. A similar pattern of upregulation was observed during the course of SARS-CoV and MERS-CoV infection of Calu3 (**Figure 3**) specifically, as well as during RSV and HPIV3 infection of A549 (**Figure 6**). Together, these results suggest that a common pattern of response to ssRNA viral infection by IFN-β and OAS-RNaseL pathway is conserved across multiple different cell lines and tissues. However, there were also several cell lines that did not display significant upregulation of *IFNB1* or genes related to the OAS-RNaseL pathway upon infection with SARS-CoV-2. Some other viruses like IAV also failed to induce these genes after infecting certain cell types like A549, despite also being ssRNA viruses (**Figure 6**). This suggests that not every cell type and virus is capable of inducing high levels of interferon and OAS under every circumstance. Upon analyzing correlations between ACE2 expression and OAS-RNaseL pathway stimulation, it was determined that the ACE2 base mean generally correlated with IFN-β and the OAS-RNaseL pathway induction following SARS-CoV-2 infection (**Figure 4**). This indicates that entry receptor abundance may be responsible for differences in OAS-RNaseL pathway response patterns. However, further studies establishing any causative relationship will need to be completed to better understand the connection.

### The Extent of OAS Pathway Upregulation in SARS-CoV-2 Appears to be Cell Line Dependent

Previous studies have found that SARS-CoV-2 replicates efficiently in Calu3 but not in A549, which could be a consequence of differences in ACE2 base mean and potentially explain why we observed strong upregulation of IFN-β and OAS-RNaseL pathway genes in Calu3 but not A549 following SARS-CoV-2 infection^32^ (**Table 1**). Caco2 was found to support replication of SARS-CoV-2 and demonstrated induction of IFN-β or OAS-RNaseL genes and had a low ACE2 base mean (**Figure 3**). While Calu3 and Caco2 were found to support similar levels of SARS-CoV-2 replication through RNAseq viral genome alignment analysis at 24 hours post infection, it cannot be ruled out that OAS-RNaseL pathway induction plays a role in inhibiting SARS-CoV-2 replication in Calu3^30^. Based on a preliminary literature review summarized in **Table 1**, it is possible that Calu3 may support significantly higher levels of SARS-CoV-2 replication in the absence of OAS-RNaseL pathway induction. In fact, the discovery that interferon signaling inhibition increases SARS-CoV-2 replication in Calu3 makes this a likely outcome^33^. However, further experimentation will need to be conducted to determine the exact role of the OAS-RNaseL pathway among all ISGs. The presence of a detectable level of viral replication despite induction of OAS-RNaseL, however, is indicative that the pathway alone is insufficient for prevention of viral replication and other parallel immune mechanisms are needed. Additionally, differentially induced innate immune mechanisms across various cell lines could also affect SARS-CoV-2 replication. It is important to note at this time that, given the intercellular connectivity and cell extrinsic aspect of immune response, these statements are preliminary and require further research to corroborate. That being said, these reasons may justify why hiPSC-CM, despite displaying strong OAS upregulation, is capable of sustaining moderate levels of viral replication. In comparison to Caco2, HT-29 cells, which are also derived from colorectal tissue, conversely demonstrated strong OAS-RNaseL pathway induction patterns more similar to that of Calu3 while also expressing a high ACE2 base mean based on our analysis (**Figure 3**). Newfound research has shown that HT-29 cells failed to support SARS-CoV-2 replication to the same degree as Caco2^34^.

### The OAS Pathway Is an Effective Antiviral Mechanism in Many ssRNA Viruses

Similar to previous findings with SeV, a virus known to induce prolific expression of interferons and OAS^35^, our analysis indicated that RNA viruses like SARS-CoV, MERS-CoV, RSV and HPIV3 were able to induce the OAS-RNaseL pathway in a similar way as SARS-CoV-2 in the same cell cultures that saw significant induction when infected with SARS-CoV-2 (**Figures 5-6**). This is consistent with known activation mechanisms of the pathway and the activity of its effectors. Differences in tropism observed for SARS-CoV-2 were also seen in SARS-CoV, which similarly failed to induce OAS-RNaseL pathway genes in H1299 and Caco2 (**Figure 5**). This preliminary data suggests a potential area of further study to determine the similarities in OAS-RNaseL pathway induction across other viruses related to SARS-CoV-2.

The importance of the OAS-RNaseL pathway in controlling viral infection was demonstrated when its inhibition led to a significant increase of viral titer in human airway epithelial cells^9,36^. *OASL*, in particular, was implicated in anti-RSV response^9,36^. Muted *OAS1* induction in airway epithelial cells of cystic fibrosis patients also correlated with increased HPIV3 replication relative to normal cells, suggesting that, at the very least, OAS1 expression is also important in controlling HPIV3^37^. Constitutive *OAS1* expression in several different contexts was also found to reduce titers of encephalomyocarditis virus (EMCV), mengovirus, and dengue virus (DENV), although direct causation is not completely understood in these contexts^38–41^. All of the previously mentioned viruses described are single-stranded RNA viruses similar to SARS-CoV-2, which suggests that the OAS-RNaseL pathway could likely play an important role in launching antiviral response against SARS-CoV-2.

### Upregulation of the OAS-RNaseL Pathway Could Promote Essential Immune Mechanisms Such as Apoptosis but Its Effect on Viral Replication Remains Unclear

Previously analyzed cell lines and tissue types that were found to strongly induce OAS-RNaseL pathway induction upon SARS-CoV-2 infection generally resulted in better outcomes following infection. SARS-CoV-2-infected Caco2 cells, which displayed weak OAS upregulation, remained viable for up to 120 hours post-infection^32^, while infected Calu3 cells, which exhibited strong OAS upregulation, displayed evidence of apoptosis^32,42^. RNaseL degradation of host rRNA and viral RNA has been found to be a contributing factor to apoptosis induction^43^. This could justify why SARS-CoV-2-infected cell lines with robust OAS-RNaseL pathway upregulation displayed significant apoptosis. However, among all cell lines analyzed, we did not find a relationship between OAS-RNaseL pathway induction and SARS-CoV-2 replication (**Figure 3**)(**Table 1**). The limitations of comparing different cell lines, such as not taking into account differing viral tropism, likely contributed to this. Further studies through direct experimentation would be necessary to gain a more complete picture of the relationship between the OAS-RNaseL mechanism and the ability for SARS-CoV-2 to replicate. Recent studies, involving the use of RNaseL knockout strains of A549-ACE2, have revealed that the deletion of RNaseL results in significantly increased SARS-CoV-2 replication, which is a promising development^44^. Combined with the clinical finding that increased *OAS1* expression positively correlated with better COVID-19 outcomes^45^, the ability of the OAS-RNaseL pathway to induce protective immune mechanisms such as apoptosis make it an attractive target for continued study as a SARS-CoV-2 therapeutic.

### Entry Receptor Abundance and the Presence of Antiviral Proteins Influence IFN Expression Pattern

One question that arises from our results relates to the expression pattern of type I and II IFNs, which have been shown to stimulate the expression of antiviral effectors including the OAS-RNaseL pathway^7,8^. We observed significant differences in induction strength for type I and II IFNs as well as associated ISGs across the different analyzed cell lines. Previous studies have also found divergent IFN induction levels between lung and intestinal epithelial cells in the context of SARS-CoV-2, though the causation is currently unknown^46^. Our analysis of ACE2 entry receptor expression levels in our conditions revealed a seeming correlation with IFN and OAS induction. This suggests that the ability of SARS-CoV-2 to attach and enter a cell could potentially influence the ability for the IFN pathway to be induced. However, additional studies are required to further confirm this relationship and prove causation. Entry receptor base means from RSV (IGF-1R, base mean = 496.8) also correlated with IFN and ISG induction in A549 cells, further corroborating this hypothesis^47^. However, we also found that two viruses using identical entry receptors, IAV and HPIV3, exhibit divergent induction patterns of IFN and ISGs, suggesting that entry receptor expression alone does not explain induction patterns. One possible explanation of divergent OAS-RNaseL responses could be based on the various non-structural viral proteins with anti-IFN activities found in SARS-CoV-2, which may play a role in modulating IFN responses in infected cells^48,49^. Similar proteins have also been documented in other RNA viruses. Knockout strains of the viral protein DS1 have demonstrated to induce significantly stronger IFN responses in NHBE compared to wild type strains^27^. SARS-CoV-2, while known to be able to induce IFN responses, was found to induce significantly lower levels of IFN-β mRNA in type II alveolar cells and cardiomyocytes compared to the Sindbis virus, which is known to strongly induce dsRNA activated pathways^44^. This would indicate that, despite the significant induction of IFN pathways seen in our results, such levels could be attenuated through the actions of anti-IFN viral proteins.

### IFN Therapy for COVID-19 Shown to be Most Effective Several Days After Symptom Onset and Could Elicit Adverse Reactions

Though treatment with IFN may seem like a suitable therapeutic option, there is potential for serious adverse events and even a lack of efficacy due to patient immune responses. One study showed that upon treatment with IFN-β for patients with relapsing-remitting multiple sclerosis, 13 of 40 patients had autoantibodies present and exhibited alterations in thyroid and liver function^50^. A recent clinical study has shown that approximately 14% of patients with severe COVID-19 induced pneumonia were confirmed to possess autoantibodies in their blood, at minimum, against one type I IFN^51^. 2.6% of women and 12.5% of men are demonstrated to be at risk for severe COVID-19 pneumonia due to autoantibodies for type I IFN^51^. That being said, a treatment regimen of IFN-β combined with lopinavir-ritonavir was found to be effective in reducing mortality in hospitalized MERS patients, though treatment was best when started seven days after symptom onset^52^. Given the longer incubation period of SARS-CoV-2, this treatment protocol may not be tenable, and as such, other therapeutic options ought to be explored.

### OAS Pathway Genes Appear to be Upregulated Independently of IFN in Epithelial Cell Lines

Another noteworthy pattern observed was the upregulation of OAS genes in the absence of *IFNB1* upregulation, which was seen in SARS-CoV-2 infected epithelial cell lines like A549 and NHBE (**Figure 7**). Type III IFNs, which are the predominant IFNs induced in epithelial cells^53,54^, were examined to determine whether or not they played a significant role in OAS activation. Despite this, our analysis showed that there was no significant upregulation or downregulation of type III IFNs in SARS-CoV-2-infected NHBE and A549 cell lines. The expression pattern of *IFNG*, the sole member of the type II IFN group, was also assessed, and did not display significant upregulation or downregulation either (**Figure 7**). One reason for this lack of IFN activation could be the result of a host-protection mechanism. For example, prolonged exposure to IFNs in IAV-infected primary murine epithelial cells was found to disrupt epithelial cell repair and proliferation^55^.

IFN has been discouraged as a therapeutic option due to the possibility of overstimulating the cytokine response^26^. Additionally, a recent study showed that 14% of patients with severe COVID-19 pneumonia were found to have autoantibodies against two forms of type I IFN proteins^51^. These outcomes demonstrate that IFN therapy may be unsuitable for severe COVID-19 cases and, as such, other treatment options ought to be explored.

### IFN-Independent OAS Pathway Activation Is a Promising Form of Therapy for Severe COVID-19 Cases

Few studies have been conducted to determine the process by which the OAS-RNaseL pathway can be activated independently of IFN. However, existing literature has shown striking results. Evidence from Kristiansen *et al.* demonstrated that exogenous OAS1 has antiviral effects independent of the type I IFN pathway, a pathway whose positive activity is directly correlated to OAS upregulation^28^. OAS was believed to be inherently linked to IFN and was thus disregarded as a viable option for antiviral drug development. However, in addition to evading IFN upregulation, the alternate OAS pathway also averts RNaseL activation^19^. This could reduce harmful side effects to the host organism since RNaseL production by healthy cells, as stimulated by paracrine signaling by IFNs, can degrade host RNA^56^. Additionally, Zhu *et al.* found that IAV is unable to evade antiviral activity by OASL^57^. However, there is data from RSV infection that shows that treatment with OASL is not as effective as Zhu *et al.* demonstrated for IAV, suggesting a viral-specific therapeutic efficacy by OASL^36^. Treatments acting independently of RNaseL and IFN upregulation while still exhibiting potent antiviral effects ought to be explored. Future research investigating the molecular details of the alternative OAS pathway could elucidate its antiviral mechanism, provide evidence for or against the therapeutic capacity of OAS1 and OASL against SARS-CoV-2 and similar viruses.

## References

1. COVID-19 Map - Johns Hopkins Coronavirus Resource Center. Accessed April 17, 2021. https://coronavirus.jhu.edu/map.html

2. Coronaviridae Study Group of the International Committee on Taxonomy of Viruses. The species Severe acute respiratory syndrome-related coronavirus: classifying 2019-nCoV and naming it SARS-CoV-2. Nat Microbiol. 2020;5(4):536–544. doi:10.1038/s41564-020-0695-z

3. Gaunt ER, Hardie A, Claas ECJ, Simmonds P, Templeton KE. Epidemiology and Clinical Presentations of the Four Human Coronaviruses 229E, HKU1, NL63, and OC43 Detected over 3 Years Using a Novel Multiplex Real-Time PCR Method. J Clin Microbiol. 2010;48(8):2940–2947. doi:10.1128/JCM.00636-10

4. Cui J, Li F, Shi Z-L. Origin and evolution of pathogenic coronaviruses. Nat Rev Microbiol. 2019;17(3):181–192. doi:10.1038/s41579-018-0118-9

5. Chen N, Zhou M, Dong X, et al. Epidemiological and clinical characteristics of 99 cases of 2019 novel coronavirus pneumonia in Wuhan, China: a descriptive study. The Lancet. 2020;395(10223):507–513. doi:10.1016/S0140-6736(20)30211-7

6. Long Q-X, Tang X-J, Shi Q-L, et al. Clinical and immunological assessment of asymptomatic SARS-CoV-2 infections. Nat Med. 2020;26(8):1200–1204. doi:10.1038/s41591-020-0965-6

7. Park A, Iwasaki A. Type I and Type III Interferons – Induction, Signaling, Evasion, and Application to Combat COVID-19. Cell Host Microbe. 2020;27(6):870–878. doi:10.1016/j.chom.2020.05.008

8. Sadler AJ, Williams BRG. Interferon-inducible antiviral effectors. Nat Rev Immunol. 2008;8(7):559–568. doi:10.1038/nri2314

9. Silverman RH. Viral Encounters with 2’,5’-Oligoadenylate Synthetase and RNase L during the Interferon Antiviral Response. J VIROL. 2007;81:10.

10. Behera AK, Kumar M, Lockey RF, Mohapatra SS. 2’-5’ Oligoadenylate Synthetase Plays a Critical Role in Interferon-γ Inhibition of Respiratory Syncytial Virus Infection of Human Epithelial Cells. J Biol Chem. 2002;277(28):25601–25608. doi:10.1074/jbc.M200211200

11. Samuel MA, Whitby K, Keller BC, et al. PKR and RNase L Contribute to Protection against Lethal West Nile Virus Infection by Controlling Early Viral Spread in the Periphery and Replication in Neurons. J Virol. 2006;80(14):7009–7019. doi:10.1128/JVI.00489-06

12. Han J-Q, Barton DJ. Activation and evasion of the antiviral 2’-5’ oligoadenylate synthetase/ribonuclease L pathway by hepatitis C virus mRNA. RNA. 2002;8(4):512–525. doi:10.1017/S1355838202020617

13. Maitra RK, Silverman RH. Regulation of Human Immunodeficiency Virus Replication by 2’,5’-Oligoadenylate-Dependent RNase L. J Virol. 1998;72(2):1146–1152. doi:10.1128/JVI.72.2.1146-1152.1998

14. Thornbrough JM, Jha BK, Yount B, et al. Middle East Respiratory Syndrome Coronavirus NS4b Protein Inhibits Host RNase L Activation. 2016;7(2):12.

15. Nogales A, Martinez-Sobrido L, Topham D, DeDiego M. Modulation of Innate Immune Responses by the Influenza A NS1 and PA-X Proteins. Viruses. 2018;10(12):708. doi:10.3390/v10120708

16. Austin BA, James C, Silverman RH, Carr DJJ. Critical Role for the Oligoadenylate Synthetase/RNase L Pathway in Response to IFN-β during Acute Ocular Herpes Simplex Virus Type 1 Infection. J Immunol. 2005;175(2):1100–1106. doi:10.4049/jimmunol.175.2.1100

17. Eberle KC, McGill JL, Reinhardt TA, Sacco RE. Parainfluenza Virus 3 Blocks Antiviral Mediators Downstream of the Interferon Lambda Receptor by Modulating Stat1 Phosphorylation. Perlman S, ed. J Virol. 2016;90(6):2948–2958. doi:10.1128/JVI.02502-15

18. Bisbal C, Martinand C, Silhol M, Lebleu B, Salehzada T. Cloning and Characterization of a RNase L Inhibitor.: A NEW COMPONENT OF THE INTERFERON-REGULATED 2-5A PATHWAY. J Biol Chem. 1995;270(22):13308–13317. doi:10.1074/jbc.270.22.13308

19. Gusho E, Zhang R, Jha BK, et al. Murine AKAP7 Has a 2’,5’-Phosphodiesterase Domain That Can Complement an Inactive Murine Coronavirus ns2 Gene. Virgin H, ed. mBio. 2014;5(4):e01312–14. doi:10.1128/mBio.01312-14

20. Kubota K, Nakahara K, Ohtsuka T, et al. Identification of 2’-Phosphodiesterase, Which Plays a Role in the 2-5A System Regulated by Interferon. J Biol Chem. 2004;279(36):37832–37841. doi:10.1074/jbc.M400089200

21. Poulsen JB, Andersen KR, Kjær KH, Vestergaard AL, Justesen J, Martensen PM. Characterization of human phosphodiesterase 12 and identification of a novel 2’-5’ oligoadenylate nuclease – The ectonucleotide pyrophosphatase/phosphodiesterase 1. Biochimie. 2012;94(5):1098–1107. doi:10.1016/j.biochi.2012.01.012

22. Lau SKP, Lau CCY, Chan K-H, et al. Delayed induction of proinflammatory cytokines and suppression of innate antiviral response by the novel Middle East respiratory syndrome coronavirus: implications for pathogenesis and treatment. J Gen Virol. 2013;94(12):2679–2690. doi:10.1099/vir.0.055533-0

23. Yoshikawa T, Hill TE, Yoshikawa N, et al. Dynamic Innate Immune Responses of Human Bronchial Epithelial Cells to Severe Acute Respiratory Syndrome-Associated Coronavirus Infection. Morty RE, ed. PLoS ONE. 2010;5(1):e8729. doi:10.1371/journal.pone.0008729

24. Cameron MJ, Ran L, Xu L, et al. Interferon-Mediated Immunopathological Events Are Associated with Atypical Innate and Adaptive Immune Responses in Patients with Severe Acute Respiratory Syndrome. J Virol. 2007;81(16):8692–8706. doi:10.1128/JVI.00527-07

25. Chan JF-W, Yao Y, Yeung M-L, et al. Treatment With Lopinavir/Ritonavir or Interferon-β1b Improves Outcome of MERS-CoV Infection in a Nonhuman Primate Model of Common Marmoset. J Infect Dis. 2015;212(12):1904–1913. doi:10.1093/infdis/jiv392

26. Sallard E, Lescure F-X, Yazdanpanah Y, Mentre F, Peiffer-Smadja N. Type 1 interferons as a potential treatment against COVID-19. Antiviral Res. 2020;178:104791. doi:10.1016/j.antiviral.2020.104791

27. Blanco-Melo D, Nilsson-Payant BE, Liu W-C, et al. Imbalanced Host Response to SARS-CoV-2 Drives Development of COVID-19. Cell. 2020;181(5):1036–1045.e9. doi:10.1016/j.cell.2020.04.026

28. Kristiansen H, Scherer CA, McVean M, et al. Extracellular 2’-5’ Oligoadenylate Synthetase Stimulates RNase L-Independent Antiviral Activity: a Novel Mechanism of Virus-Induced Innate Immunity. J Virol. 2010;84(22):11898–11904. doi:10.1128/JVI.01003-10

29. Afgan E, Baker D, Batut B, et al. The Galaxy platform for accessible, reproducible and collaborative biomedical analyses: 2018 update. Nucleic Acids Res. 2018;46(W1):W537–W544. doi:10.1093/nar/gky379

30. Cao Y, Xu X, Kitanovski S, et al. Comprehensive Comparison of Transcriptomes in SARS-CoV-2 Infection: Alternative Entry Routes and Innate Immune Responses. Bioinformatics; 2021. doi:10.1101/2021.01.07.425716

31. Sharma A, Garcia G, Wang Y, et al. Human iPSC-Derived Cardiomyocytes Are Susceptible to SARS-CoV-2 Infection. Cell Rep Med. 2020;1(4):100052. doi:10.1016/j.xcrm.2020.100052

32. Chu H, Chan JF-W, Yuen TT-T, et al. Comparative tropism, replication kinetics, and cell damage profiling of SARS-CoV-2 and SARS-CoV with implications for clinical manifestations, transmissibility, and laboratory studies of COVID-19: an observational study. Lancet Microbe. 2020;1(1):e14–e23. doi:10.1016/S2666-5247(20)30004-5

33. Schroeder S, Pott F, Niemeyer D, et al. Interferon antagonism by SARS-CoV-2: a functional study using reverse genetics. Lancet Microbe. Published online March 2021:S2666524721000276. doi:10.1016/S2666-5247(21)00027-6

34. Bojkova D, McGreig JE, McLaughlin K-M, et al. SARS-CoV-2 and SARS-CoV Differ in Their Cell Tropism and Drug Sensitivity Profiles. Microbiology; 2020. doi:10.1101/2020.04.03.024257

35. Melchjorsen J, Kristiansen H, Christiansen R, et al. Differential Regulation of the OASL and OAS1 Genes in Response to Viral Infections. J Interferon Cytokine Res. 2009;29(4):199–208. doi:10.1089/jir.2008.0050

36. Dhar J, Cuevas RA, Goswami R, Zhu J, Sarkar SN, Barik S. 2’-5’-Oligoadenylate Synthetase-Like Protein Inhibits Respiratory Syncytial Virus Replication and Is Targeted by the Viral Nonstructural Protein 1. Dermody TS, ed. J Virol. 2015;89(19):10115–10119. doi:10.1128/JVI.01076-15

37. Zheng S, De BP, Choudhary S, et al. Impaired Innate Host Defense Causes Susceptibility to Respiratory Virus Infections in Cystic Fibrosis. Immunity. 2003;18(5):619–630. doi:10.1016/S1074-7613(03)00114-6

38. Coccia EM, Albertini R, Angela I, Rossi B, Chebath J. A Full-Length Murine 2-5A Synthetase cDNA Transfected in NIH-3T3 Cells Impairs EMCV but Not VSV Replication. Virology. 1990;179:6.

39. Rysiecki G, Gewert DR, Williams BRG. Constitutive Expression of a 2’,5’-Oligoadenylate Synthetase cDNA Results in Increased Antiviral Activity and Growth Suppression. J Interferon Res. 1989;9(6):649–657. doi:10.1089/jir.1989.9.649

40. Chebath J, Benech P, Revel M, Vigneron M. Constitutive expression of (2’-5’) oligo A synthetase confers resistance to picornavirus infection. Lett Nat. 1987;330:2.

41. Lin R-J, Yu H-P, Chang B-L, Tang W-C, Liao C-L, Lin Y-L. Distinct Antiviral Roles for Human 2’,5’-Oligoadenylate Synthetase Family Members against Dengue Virus Infection. J Immunol. 2009;183(12):8035–8043. doi:10.4049/jimmunol.0902728

42. Li S, Zhang Y, Guan Z, et al. SARS-CoV-2 triggers inflammatory responses and cell death through caspase-8 activation. Signal Transduct Target Ther. 2020;5(1):235. doi: 10.1038/s41392-020-00334-0

43. Castelli JC, Hassel BA, Maran A, et al. The role of 2’-5’ oligoadenylate-activated ribonuclease L in apoptosis. Cell Death Differ. 1998;5(4):313–320. doi:10.1038/sj.cdd.4400352

44. Li Y, Renner DM, Comar CE, et al. SARS-CoV-2 Induces Double-Stranded RNA-Mediated Innate Immune Responses in Respiratory Epithelial Derived Cells and Cardiomyocytes. Microbiology; 2020. doi:10.1101/2020.09.24.312553

45. Zhou S, Butler-Laporte G, Nakanishi T, et al. Circulating Proteins Influencing COVID-19 Susceptibility and Severity: a Mendelian Randomization Study.:27.

46. Stanifer ML, Kee C, Cortese M, et al. Critical Role of Type III Interferon in Controlling SARS-CoV-2 Infection in Human Intestinal Epithelial Cells. Cell Rep. 2020;32(1):107863. doi: 10.1016/j.celrep.2020.107863

47. Griffiths CD, Bilawchuk LM, McDonough JE, et al. IGF1R is an entry receptor for respiratory syncytial virus. Nature. 2020;583(7817):615–619. doi:10.1038/s41586-020-2369-7

48. Xia H, Cao Z, Xie X, et al. Evasion of Type I Interferon by SARS-CoV-2. Cell Rep. 2020;33(1):108234. doi:10.1016/j.celrep.2020.108234

49. Lei X, Dong X, Ma R, et al. Activation and evasion of type I interferon responses by SARS-CoV-2. Nat Commun. 2020;11(1):3810. doi:10.1038/s41467-020-17665-9

50. Durelli L, Ferrero B, Oggero A, et al. Autoimmune events during interferon beta-1b treatment for multiple sclerosis. J Neurol Sci. 1999;162(1):74–83. doi:10.1016/S0022-510X(98)00299-8

51. Bastard P, Rosen LB, Zhang Q, et al. Autoantibodies against type I IFNs in patients with life-threatening COVID-19. Science. 2020;370(6515). doi:10.1126/science.abd4585

52. Arabi YM, Asiri AY, Assiri AM, et al. Interferon Beta-1b and Lopinavir–Ritonavir for Middle East Respiratory Syndrome. N Engl J Med. Published online 2020:12.

53. Reid E, Charleston B. Type I and III Interferon Production in Response to RNA Viruses. J Interferon Cytokine Res. 2014;34(9):649–658. doi:10.1089/jir.2014.0066

54. Okabayashi T, Kojima T, Masaki T, et al. Type-III interferon, not type-I, is the predominant interferon induced by respiratory viruses in nasal epithelial cells. Virus Res. 2011;160(1-2):360–366. doi:10.1016/j.virusres.2011.07.011

55. Major J, Crotta S, Llorian M, et al. Type I and III interferons disrupt lung epithelial repair during recovery from viral infection. Science. 2020;369(6504):712–717. doi:10.1126/science.abc2061

56. Karasik A, Jones GD, DePass AV, Guydosh NR. NAR Breakthrough Article Activation of the antiviral factor RNase L triggers translation of non-coding mRNA sequences.:20.

57. Zhu J, Ghosh A, Sarkar SN. OASL—a new player in controlling antiviral innate immunity. Curr Opin Virol. 2015;12:15–19. doi:10.1016/j.coviro.2015.01.010

58. Emanuel W, Kirstin M, Vedran F, et al. Bulk and Single-Cell Gene Expression Profiling of SARS-CoV-2 Infected Human Cell Lines Identifies Molecular Targets for Therapeutic Intervention. Systems Biology; 2020. doi:10.1101/2020.05.05.079194

59. Yuan S, Chu H, Chan JF-W, et al. SREBP-dependent lipidomic reprogramming as a broad-spectrum antiviral target. Nat Commun. 2019;10(1):120. doi:10.1038/s41467-018-08015-x

60. Guo Y, Luo R, Wang Y, et al. *Modeling SARS-CoV-2 Infection* in Vitro *with a Human Intestine-on-Chip Device*. Bioengineering; 2020. doi:10.1101/2020.09.01.277780

