## Supplementary table 1 for "Comparing Expression of OAS-RNaseL Pathway-Related Genes in SARS-CoV-2 and Similar Viruses"

### Supplemental Materials

**Supplemental Table 1.** Summary of publicly available BioProjects and SRA run accessions used in this study.

| BioProject Accession | Cell line or sample | Virus or treatment | SRA run accessions |
|---|---|---|---|
| *PRJNA625518* | Calu3 lung epithelium adenocarcinoma cells | SARS-CoV-2 (MOI 0.33)[58] | **Mock:** SRR11549979, SRR11549980, SRR11549981, SRR11549982, **4hpi:** SRR11550009, SRR11550010 **12hpi:** SRR11550003, SRR11550004 **24hpi:** SRR11549993, SRR11549994 |
| | | SARS-CoV (MOI 0.33) | **Mock:** SRR11549979, SRR11549980, SRR11549981, SRR11549982 **4hpi:** SRR11549983, SRR11549984 **12hpi:** SRR11549985, SRR11549986 **24hpi:** SRR11549987, SRR11549988 |
| | H1299 lung lymph node large cell carcinoma | SARS-CoV-2 (MOI 0.33) | **Mock:** SRR11549939, SRR11549940 **4 hpi:** SRR11549955, SRR11549956 **12 hpi:** SRR11549949, SRR11549950 **24 hpi:** SRR11549951, SRR11549952 |
| | | SARS-CoV (MOI 0.33) | **Mock:** SRR11549939, SRR11549940 **4 hpi:** SRR11549947, SRR11549948 **12 hpi:** SRR11549941, SRR11549942 **24 hpi:** SRR11549943, SRR11549944 |
| | Caco2 colorectal epithelial adenocarcinoma cells | SARS-CoV-2 (MOI 0.33) | **Mock:** SRR11549961, SRR11549962 **4 hpi:** SRR11549973, SRR11549974 **12 hpi:** SRR11549969, SRR11549970 **24 hpi:** SRR11549971, SRR11549972 |
| | | SARS-CoV (MOI 0.33) | **Mock:** SRR11549961, SRR11549962 **4 hpi:** SRR11549967, SRR11549968 **12 hpi:** SRR11549963, SRR11549964 **24 hpi:** SRR11549965, SRR11549966 |
| *PRJNA506733* | Calu3 lung epithelium adenocarcinoma cells | MERS-CoV (MOI 2)[59] | **Mock:** SRR8239989, SRR8239990, SRR8239991 **24hpi:** SRR8239992, SRR8239993, SRR8239994 |
| *PRJNA615032* | Normal human bronchial epithelial (NHBE) | SARS-CoV-2 (MOI 2) | **Mock:** SRR11412215-26 (n=12) **24hpi:** SRR11412227-38 (n=12) |
| | | IAV (MOI 3) | **Mock:** SRR11517759-74 (n=16) **12hpi:** SRR11517775-90 (n=16) |

| | | | |
|---|---|---|---|
| | | IFN-β (100U/mL) | **Mock:** SRR11517759-74 (n=16)<br>**12hpi:** SRR11517823-30 (n=8) |
| | A549 lung epithelium carcinoma cells | SARS-CoV-2 (MOI 0.2) | **Mock:** SRR11412239-50, SRR11412263-70 (n=20)<br>**24hpi:** SRR11412251-62 (n=12) |
| | | SARS-CoV-2 (MOI 2) | **Mock:** SRR11517674-76 (n=3)<br>**24hpi:** SRR11517677-79 (n=3) |
| | | RSV (MOI 15) | **Mock:** SRR11412263-67 (n=5)<br>**24hpi:** SRR11412271-75 (n=5) |
| | | IAV (MOI 5) | **Mock:** SRR11412283-86 (n=4)<br>**9hpi:** SRR11412287-90 (n=4) |
| | | HPIV3 (MOI 3) | **Mock:** SRR11517750-52 (n=3)<br>**24hpi:** SRR11517756-58 (n=3) |
| | A549-ACE2 lung epithelium carcinoma cells containing a human ACE2 vector | SARS-CoV-2 (MOI 0.2) | **Mock:** SRR11517680-82 (n=3)<br>**24hpi:** SRR11517741-43 (n=3) |
| | | SARS-CoV-2 (MOI 2) | **Mock:** SRR11573892-97 (n=6)<br>**24hpi:** SRR11573904-09 (n=6) |
| | Lung biopsy | SARS-CoV-2 | **Heated negative control:** SRR11517725-32 (n=8)<br>**COVID-19 postmortem:** SRR11517733-40 (n=8) |
| *PRJNA658711* [60] | Human umbilical vein endothelial cells (HUVEC) | SARS-CoV-2 | **Mock:** SRR12495888, SRR12495887, SRR12495886<br>**Infected:** SRR12495885, SRR12495893, SRR12495894 |
| | HT-29 colon epithelial cells | SARS-CoV-2 | **Mock:** SRR12495896, SRR12495895, SRR12495892,<br>**Infected:** SRR12495891, SRR12495890, SRR12495889 |
| *PRJNA634489* | Platelets from COVID-19 patients under intensive care | SARS-CoV-2 | **Healthy:** SRR12113314-15, SRR12113317 (n=3)<br>**COVID-19 patients:** SRR12113306-07, SRR12113309 (n=3) |
| *PRJNA631969* | hiPSC-CM cardiomyocytes | SARS-CoV-2 (MOI 0.1) | **Mock:** SRR11777737-39 (n=3)<br>**72hpi:** SRR11777734-36 (n=3) |
| *PRJNA644625* | Primary human | SARS-CoV-2 | **Mock:** SRR12168536-38 (n=3) |

| | | | |
|---|---|---|---|
| | airway epithelial cells (pHAE) | (MOI 0.25) | **48hpi:** SRR12168542-44 (n=3) |
| *PRJNA633783* | Normal human bronchial epithelial cell organoids (hBEpC) | SARS-CoV-2 | **Healthy:** SRR11811019, SRR11811020, SRR11811021 (n=3)<br>**Infected:** SRR11811022, SRR11811023, SRR11811024 (n=3) |
| *PRJNA629752* | Peripheral blood mononuclear cells (PBMC) (Low ACE2) | SARS-CoV-2 and IAV | **Healthy:** SRR11680211, SRR11680219, SRR11680220, SRR11680225 (n=4)<br>**SARS-CoV-2:** SRR11680207, SRR11680215, SRR11680216, SRR11680221 (n=4) |
| *PRJNA633393* | Peripheral blood mononuclear cells (PBMC) (High ACE2) | SARS-CoV-2 | **Healthy:** SRR11804725-30 (n=6)<br>**Infected:** SRR11804718-24 (n=7) |
| *PRJNA509906* | Normal human dermal fibroblasts (NHDF) | HSV-1 | **Healthy:** SRR8315701-3 (n=3)<br>**Infected:** SRR8315713-15 (n=3) |
